# Mathematical Modeling of Bone Remodeling after Surgical Menopause

**DOI:** 10.1101/2025.10.19.683313

**Authors:** Anna C. Nelson, Edwina F. Yeo, Yun Zhang, Carley V. Cook, Sophie Fischer-Holzhausen, Lauryn Keeler Bruce, Pritha Dutta, Samaneh Gholami, Brenda J. Smith, Ashlee N. Ford Versypt

## Abstract

Osteoporosis is a skeletal pathology characterized by decreased bone mass and structural deterioration resulting from an imbalance in bone metabolic processes. Estrogen deficiency in postmenopausal women leads to an increased risk of osteoporosis, while women who have undergone complete oophorectomies display an even higher risk due to the sudden decrease in estrogen. Some evidence indicates that bone loss slows in the period beyond 15 years after surgery; however, there is substantial uncertainty in clinical data. To explore the effects of surgically induced menopausal transition, here we propose a mathematical model for the bone cell dynamical responses to sudden estrogen deficiency, which extends an existing model for osteoporosis due to aging and natural menopause. Using data on the key impacts observed in female mice and humans after bilateral oophorectomy, this new model considers the role of osteocytes embedded within the bone mineralized matrix in regulating osteoclastogenesis, which results in increased bone resorption after surgical menopause. The values of model parameters in natural and surgical menopause were estimated from aggregated human clinical data from existing longitudinal studies. The new model effectively captures the previously unmodeled increase in bone loss during the first 15 years post-surgical menopause and the rebound in bone mineral density in the long-term. With this model, effects of treatments on targeting osteocyte dynamics could be explored in the future.

## 1 INTRODUCTION

Bone tissue is continuously resorbed and formed through the bone remodeling process, which maintains healthy tissue and repairs micro-fractures in the skeleton. At homeostasis, the biomechanical, biochemical, and cellular mechanisms involved in remodeling of the adult skeleton maintain bone mass. However, any alteration in this complex bone turnover cycle can result in changes in bone (i.e., pathologic or anabolic) (Allen and Burr, 2014). The cellular mechanisms that determine bone health occur in functional locations called basic multicellular units (BMUs) within which cellular interactions contribute to bone tissue remodeling through a continuous cycle of activation, resorption, and formation (Robling et al., 2006). The three main types of cells that contribute to this cycle are osteoclasts, osteoblasts, and osteocytes. Osteocytes, widely recognized for their strain sensitivity, are the most abundant of these cells and signal to recruit other cells to the BMU to initiate bone resorption and formation (Creecy et al., 2021). One such signal is sclerostin, which is a Wnt inhibitor secreted by osteocytes that promotes resorption through upregulating osteoclastogenesis and inhibits formation by downregulating osteoblastogenesis (Delgado-Calle et al., 2017). Once activated by sclerostin, osteoclasts degrade the bone protein matrix and solubilize the mineral hydroxyapatite (De Maré et al., 2020; Kenkre and Bassett, 2018). Osteoblasts are activated in the BMU to initiate bone matrix formation by forming osteoid tissue, which is later mineralized into bone tissue (Kenkre and Bassett, 2018). An estimated 20% of the osteoblasts within osteoid tissue differentiate further into osteocytes. After the resorbed bone tissue is replaced through bone formation, osteocytes signal to BMU cells to slow bone formation (Kenkre and Bassett, 2018).

Perturbations of the bone remodeling process can cause an imbalance in the catabolic and anabolic activity, which can lead to bone pathologies that involve substantial bone loss. In particular, osteoporosis is a low bone density disease caused by such an imbalance and leads to increased fracture risk, resulting in significant physical and financial burden to those affected by it (Harvey et al., 2010). Osteoporosis is prevalent in postmenopausal women, with approximately 50% of women over age 50 affected by low bone mass or osteoporosis (Rachner et al., 2011; Sarafrazi et al., 2021). Estrogen-deficient bone loss has been linked to several metabolic processes in bone remodeling. The presence of estrogen prevents apoptosis of osteocytes and osteoblasts, and estrogen reduces levels of sclerostin (Mödder et al., 2010). Estrogen also reduces the impact of osteoclasts by preventing osteoclastogenesis and increasing osteoclast apoptosis (Florencio-Silva et al., 2015). When estrogen levels temporarily drop during perimenopause or are permanently low after menopause, osteoclast differentiation increases and causes more bone removal (Hsu et al., 2024); meanwhile, despite an initial increase in osteoblasts with the decline in estrogen, the bone-forming osteoblast activity is unable to match the pace of the increase in resorption, resulting in less bone formation (Jilka et al., 1998; Almeida et al., 2007; Seeman, 2013; Karlamangla et al., 2021).

In both biological sexes, the BMUs are affected by estrogens (Khosla et al., 2012). In individuals with ovaries, highly variable and declining estrogen production throughout perimenopause and menopause leads to bone loss. During late perimenopause and the early postmenopausal period, bone is lost at a rate of 2–2.4% per year in the spine and 1.2-1.7% per year in the hip (Finkelstein et al., 2008; Shieh et al., 2016). During this time, women lose approximately 25% of their trabecular bone (honeycomb-shaped bone structures) and 15% of their cortical bone (dense bone tissue on the outside of bone structures) (Finkelstein et al., 2008). This rapid bone degradation lasts about 5 to 10 years. After this period, bone is lost at a much slower rate of 0.5% annually (The North American Menopause Society, 2021).

Another cause of estrogen loss is the surgical removal of ovaries. A bilateral oophorectomy is usually performed to reduce the risk of cancer or treat non-malignant ovarian diseases, such as endometriosis or benign cysts (Cohen et al., 2012; Challberg et al., 2011; Aitken et al., 1973). This sudden onset of estrogen deficiency increases a patient’s risk for osteoporosis (Rodriguez and Shoupe, 2015). After oophorectomy, both the age of the patient at surgery and the usage of hormone replacement therapies influence the likelihood of developing osteoporosis. Oophorectomy before the age of 45 was reported to lead to an increased risk for osteoporosis; however, those who underwent an oophorectomy after age 45 had similar bone density to women with intact ovaries (Aitken et al., 1973). A review of available early postmenopause data found that the earlier menopause occurred, whether natural or surgical, the lower the resulting bone density became (Gallagher, 2007). Abnormal bone scans were identified in 71% of women who underwent a preventative bilateral oophorectomy (Cohen et al., 2012). This study did not find a difference between those who underwent surgery before or after the age of natural menopause; however, the authors pointed out that there was a large difference in follow-up ages between the two groups, indicating that the age of the woman when the bone scans were taken is also an important factor that should have been accounted for (Cohen et al., 2012). Fakkert et al. (2017) provided a systematic review of bone mineral density (BMD) following surgical menopause. They cautioned about data bias in the reported data. They concluded that while surgical menopause substantially decreases BMD, this decline becomes indistinguishable from that observed after natural menopause, once the age of natural menopause is reached (Fakkert et al., 2017). Oophrectomy-induced bone loss may be prevented with hormone replacement therapy, but many women have an aversion to taking estrogen due to perceived risk (Challberg et al., 2011). Overall, the etiology of bone loss that leads to osteoporosis due to surgical menopause needs further exploration.

The impacts of surgical menopause on mechanisms involved in bone remodeling have been explored using ovariectomized animal models, where animals undergo either an ovariectomy (OVX) procedure or a control procedure (i.e., sham operation) that mimics surgery but keeps the ovaries intact, and via *in vivo* studies from human biopsies or animal cells in culture. Several sheep, rat, and mouse *in vivo* and *in vitro* studies showed that osteocyte apoptosis increased after estrogen withdrawal (Brennan et al., 2011; Tomkinson et al., 1997, 1998; Huber et al., 2007; Emerton et al., 2010; Brennan et al., 2014), while some studies showed that the number of osteocytes were only significantly lower immediately after surgery and osteocyte cell counts increased after one month post-OVX (Florencio-Silva et al., 2018). Other studies showed that the increased osteocyte apoptosis caused by a rapid loss of estrogen increased the differentiation rate of osteoclast precursors due to released receptor activator of nuclear factor κB ligand (RANKL) (McNamara, 2021; Naqvi et al., 2020; Choi et al., 2008). While OVX experiments indicated an important change in osteocyte cell number after surgical menopause, it is still unclear how these mechanisms affect overall BMD and fracture risk.

Although estrogen plays a central role in bone health, few mathematical models in systems biology focus on the mechanisms of estrogen’s impact on bone. A recent review from our team (Cook et al., 2024) identified published mathematical models that incorporate explicit effects of estrogen in bone remodeling and postmenopausal treatments (Rattanakul et al., 2003; Schmidt et al., 2011; Post et al., 2013; Berkhout et al., 2015, 2016; Chaiya and Rattanakul, 2017; Javed et al., 2018; Jörg et al., 2022) and others with implicit estrogen effects (Lemaire et al., 2004; Lemaire and Cox, 2019; Scheiner et al., 2013, 2014; Larcher and Scheiner, 2021; Trichilo et al., 2019; Martin et al., 2019). However, none of these models consider the implications of surgical menopause on bone cell populations. The recent model by Jörg et al. (2022) includes estrogen’s effects on osteoclasts and sclerostin, includes resorption signals, and incorporates osteocyte dynamics. This model is based on realistic human time frames and includes pharmacological treatments. In particular, the mathematical model is parameterized using BMD data from patients who experienced natural menopause (Looker et al., 1998) and estimates parameters using datasets that incorporate different treatment protocols. However, the natural menopause data used to parameterize this model contains only two postmenopausal data points. Furthermore, as in the other models discussed in Cook et al. (2024), the model in Jörg et al. (2022) does not consider an abrupt decline in estrogen, which occurs in the surgical menopause scenario, nor does this model incorporate important mechanisms involved in surgical menopause, such as the impact of osteocyte death on BMD.

In this paper, we (1) aggregate BMD data sources from natural menopause patients and then reparameterize a subset of parameters in the Jörg et al. (2022) model to ensure that the mechanisms of the model reflect the broader BMD trends after natural menopause, (2) include new estrogen dynamics to describe the sudden and dramatic loss of estrogen due to surgical menopause, and (3) extend the mathematical Jörg et al. (2022) model for the dynamical responses of BMU bone cells to the case of estrogen deficiency during the surgical menopausal transition using information about the critical impacts observed in female mice and humans after removal of the ovaries. The new model considers the role of embedded osteocyte cells in regulating osteoclast differentiation and inducing enhanced bone resorption after surgical menopause. With this new model, we perform parameter exploration to determine which mechanisms are most important in capturing surgical menopause data trends. This model could be used to explore medical interventions to correct the imbalances in bone remodeling after surgical menopause in a population at higher risk for early onset of osteoporosis.

## 2 METHODS

### 2.1 Curated bone mineral density (BMD) data

While the impact of gradual estrogen decline on the bone remodeling process was investigated by Jörg et al. (2022), the dataset used to parameterize their model was sparse after menopause. To improve the accuracy of the Jörg et al. (2022) model in the natural menopause scenario and to parameterize our new surgical menopause model, we gather a larger set of published postmenopausal data. In particular, we aggregate human BMD studies that investigated the impact of different types of menopause in women without hormonal replacement. Clinical studies measure BMD in locations such as L2–L4 vertebrae in the lumbar spine and the hip (i.e., proximal femur) across different age, ethnic, and sex groups. Most BMD studies were cross-sectional where dual-energy X-ray absorptiometry scans were performed on subjects across age ranges in premenopausal or postmenopausal groups, and the data were typically reported as the average and standard deviation of BMD for the subjects (Pansini et al., 1995; Ohta et al., 2002; Hadjidakis et al., 2003; Chittacharoen et al., 1999; Looker et al., 1998; Yasui et al., 2007). We also include one longitudinal study that assessed BMD in the lumbar spine and hip across 18 months in women who underwent bilateral oophorectomy before natural menopause (Hibler et al., 2016). For consistency, we utilize BMD measurements in lumbar spine from all studies except for Looker et al. (1998), which measured proximal femur BMD, as this data was used to estimate parameters in Jörg et al. (2022). The longitudinal studies all consistently reported spine data, and only a subset reported femur BMD.

To compare BMD measurements from various datasets, each dataset’s BMD values are normalized by their respective measured values at menopause onset, and time is rescaled based on the menopause onset time, which we define as *t* = *t*_*m*_ where *t* is age in years. The rescaled time is *t* − *t*_*m*_ for time since menopause onset. We plotted the average normalized BMD measurements and standard deviations for natural menopause (Figure 1a) and surgical menopause (Figure 1b) with linear fits to data in the first 15 years postmenopause. The slopes show a 1.54% decrease in BMD per year in the first 15 years for natural menopause (Figure 1a) and a 2.03% decrease in BMD per year over the same period for surgical menopause (Figure 1b). Interestingly, the BMD measurements from Hadjidakis et al. (2003) show a rebound in BMD long after surgical menopause.

**Figure 1.**
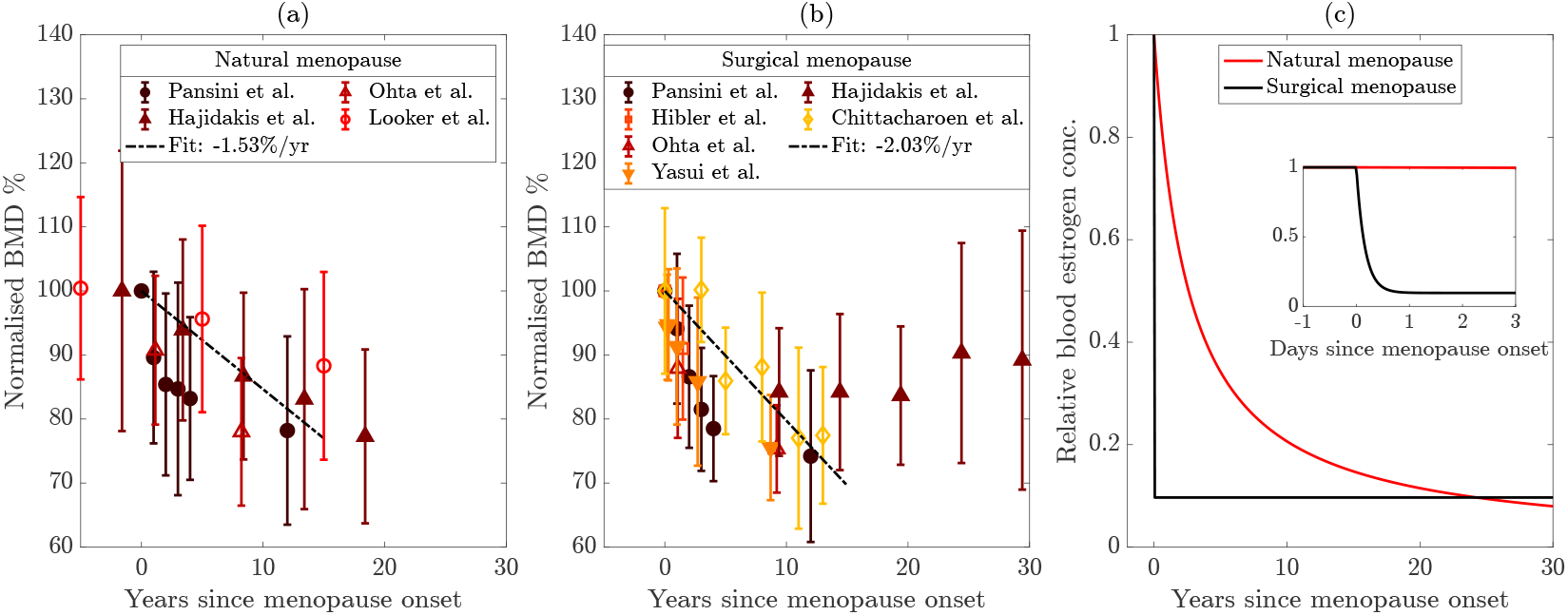
(a,b) Bone mineral density (BMD) measured in the lumbar spine of women (except Looker et al. (1998) are from the hip). Data are normalized on premenopausal levels from the time of menopause onset, and error bars show standard deviation. Normalization process for each dataset is detailed in Section 2.1. The data are from Pansini et al. (1995); Ohta et al. (2002); Hadjidakis et al. (2003); Looker et al. (1998); Hibler et al. (2016); Chittacharoen et al. (1999); Yasui et al. (2007). Dashed lines show linear fits to the data with slopes listed in the legends. (a) Natural menopause and (b) surgical menopause. (c) Comparison of estrogen decline in natural and surgical menopause, shown over 30 years after menopause onset. The inset shows the decline over the 3 days following menopause onset.

### 2.2 Modeling bone remodeling after natural and surgical menopause

To understand how natural and surgical menopause differentially impact the bone remodeling system, we utilize a mathematical framework that assumes well-mixed chemical and cellular species within a BMU. With this assumption, dynamics of the bone cell populations, chemical signals, and hormones are described with ordinary differential equations (ODEs). These species ultimately impact the formation of bone through changes in the bone production and bone degradation rates. A number of estrogen effects were included in the Jörg et al. (2022) model: the inhibition of sclerostin production (Mödder et al., 2010) and the downregulation of osteoclastic bone resorption via suppressing osteoclast differentiation (Kameda et al., 1997). Therefore, a decrease in estrogen through natural or surgical menopause increases osteoclast and sclerostin levels, which both lead to increased bone reabsorption. We now describe our model that extends the work of Jörg et al. (2022) to consider surgical menopause. To explore the impact of surgical menopause on bone remodeling, we incorporate two new mechanisms into the model: increased apoptosis of osteocytes through a term *η*(*t*) and increased differentiation of osteoclasts through a term *ω*(*t*). We include these effects through apoptosis and differentiation rates, which depend on the time since surgical menopause. This time-dependence captures inflammation and metabolic responses outside of the BMU. In Figure 2, we summarize the model in a schematic, illustrating how different cells and chemical signals interact and impact bone formation. The mechanisms impacted by the presence of estrogen are shown in Figure 2, with new surgical menopause effects specifically highlighted with red dashed arrows marked with scissor icons, representing surgical removal of the ovaries.

**Figure 2.**
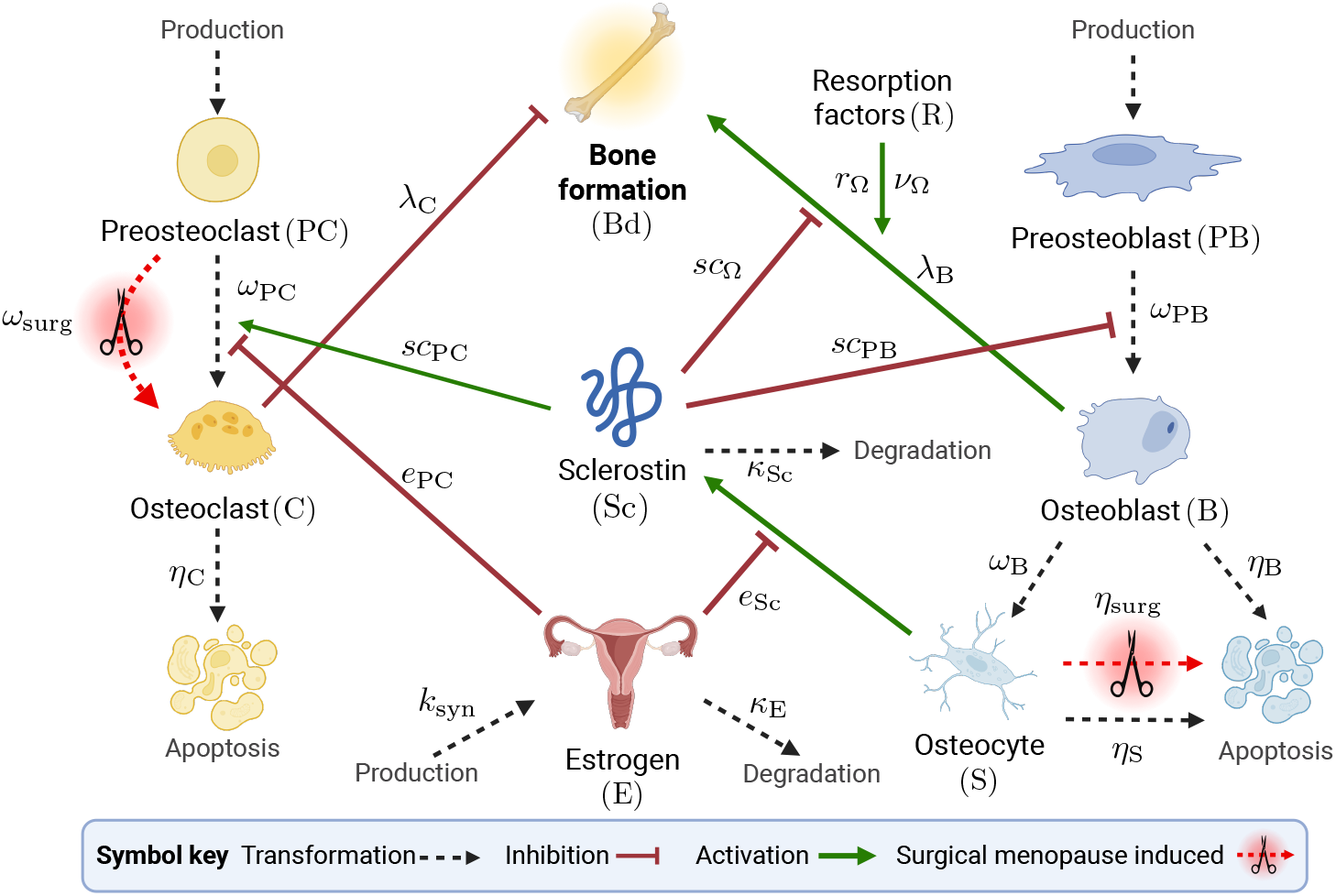
Schematic of mathematical model species in bone remodeling process. The transformations of cells through differentiation, production, and degradation are illustrated by black dashed arrows. Signaling-related inhibition interactions are shown by red flat-head arrows, and activation interactions are shown as green solid arrows. Parameters that govern interactions are shown near the corresponding arrows. The presence of estrogen is shown in the schematic as an inhibitor of osteoclast differentiation and sclerostin production. Surgical menopause-induced changes are depicted by red dashed arrows with scissors icons. Created with BioRender.com.

Each cell population and chemical concentration is scaled by reference values for ease of computation: PC and C are scaled by the number of preosteoclasts produced per day in the BMU; PB, B, S, and Sc are scaled relative to the number of preosteoblasts produced per day; and estrogen is scaled by the initial concentration of estrogen at the onset of menopause.

Activation and repression signaling interactions are modeled with saturating Hill-type functions:

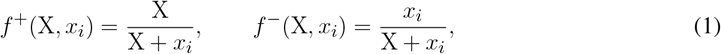

respectively. For each type of interaction by species X, the threshold concentration where half of the interaction strength is achieved is the parameter *x*_*i*_, where *i* denotes the target of the signaling interaction.

For simplicity, we assume estrogen is described by an algebraic equation, and its form depends on the type of menopause investigated. The normalized estrogen concentration over time during natural menopause is (Jörg et al., 2022):

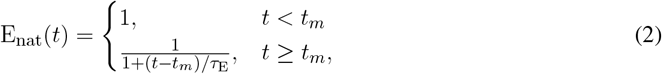

where *t*_*m*_ is the time of onset of estrogen decline and *τ*_E_ is the characteristic time of estrogen decline. We capture the sudden and rapid decrease in relative estrogen concentration due to oophorectomy surgery using the following equation, which is derived assuming estrogen is still synthesized at a reduced constant (zero-order) rate of *k*_syn_ post-surgery and degraded at a first-order rate with rate constant *κ*_E_:

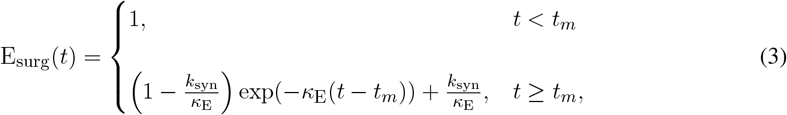

where the degradation rate of estrogen is defined as *κ*_E_ = ln(2)*/t*1*/*2 and *t*1*/*2 is the half-life of estrogen, which is 161 minutes in postmenopausal women (Ginsburg et al., 1998). The initial concentration of estrogen before surgery is 156 pg/mL, and the estrogen level stabilizes and reaches a steady state by about 30 days post-surgery to an average value of 15 pg/mL (Bellanti et al., 2013). Thus, we set the post-surgery normalized estrogen concentration as 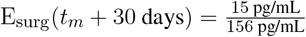. The synthesis rate after surgery is calculated as *k*_syn_ = *κ*_E_E_surg_(*t*_*m*_ + 30 days). Figure 1c shows the difference in the estrogen decrease in the case of natural menopause compared to surgical menopause, described by Eqs. (2) and (3), where the inset illustrates the timescale of rapid estrogen decline in surgical menopause.

The changes in cell number over time of the preosteoclasts (PC) and osteoclasts (C) are given by

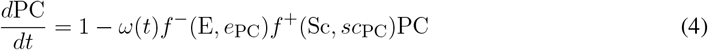

and

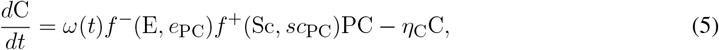

respectively. Preosteoclasts are produced at a constant basal rate of one upon scaling and differentiate into osteoclasts at a rate of *ω*(*t*). The presence of estrogen inhibits osteoclast differentiation, while sclerostin activates this differentiation with thresholds *e*_PC_ and *sc*_PC_, respectively. Osteoclast apoptosis occurs at a rate *η*_C_. Note that we do not include the apoptosis of any precursor cells (preosteoclasts or preosteoblasts) or any effect of estrogen on osteoclast apoptosis as these were estimated to have a negligible impact in Jörg et al. (2022).

The first effect of surgical menopause is included in a new time-dependent differentiation rate for preosteoclasts to osteoclasts, *ω*(*t*), defined as

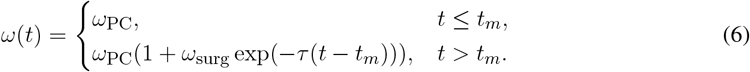

Before estrogen decline *t* ≤ *t*_*m*_ and for natural menopause according to the Jörg et al. (2022) model, differentiation occurs with a constant rate *ω*_PC_. At the onset of rapidly decreased estrogen due to surgical menopause (Eq. (3) and Figure 1c), the differentiation rate increases by *ω*_surg_, e.g., for a 10% increase, then *ω*_surg_ = 0.1. This increased rate of differentiation lasts for a period defined by the parameter *τ*, such that longer-lasting effects of surgery are defined by smaller *τ* .

The preosteoblast (PB) and osteoblasts (B) populations are governed by

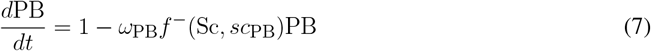

and

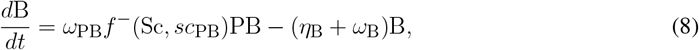

respectively. Preosteoblasts are produced at a constant basal rate of one upon scaling and differentiate into osteoblasts at a rate of *ω*_PB_ inhibited by sclerostin. Osteoblasts have an apoptosis rate of *η*_B_ and are further differentiated into osteocytes at a rate of *ω*_B_.

The dynamic population of osteocytes (S) is described by

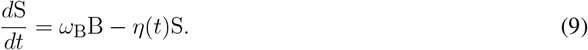

Osteocytes are derived from osteoblasts and are removed at a rate dependent on *η*(*t*).

The second effect of surgical menopause is increased apoptosis of osteocytes via the new time-dependent rate, *η*(*t*), defined as

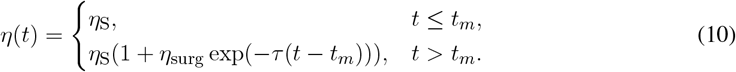

Similarly to Eq. (6), we assume that before menopause and for natural menopause according to the Jörg et al. (2022) model, differentiation occurs at a constant rate of *η*_S_. At surgical menopause onset, apoptosis increases by *η*_surg_, then returns to previous levels over a timescale of *τ* . This timescale is assumed to be equal to the timescale of increased osteoclast differentiation in Eq. (6).

The production of the signaling molecule sclerostin is governed by

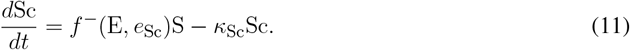

where sclerostin is produced by osteocytes at a rate inhibited by estrogen with threshold *e*_Sc_ and is degraded at rate *κ*_Sc_. Sclerostin affects bone formation through activation of osteoclastogenesis in Eqs. (4) and (5) and inhibition of osteoblastogenesis in Eqs. (7) and (8), respectively.

Bone density, Bd, is determined by

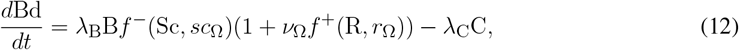

where the resorption factor R is assumed to be equal to the amount of osteoclasts present, i.e., R = C. Osteoclasts inhibit bone formation, and osteoblasts contribute to bone formation. From Eq. (12), the rate of bone density change increases proportionally to osteoblast number at a rate of *λ*_B_. This production is modulated by sclerostin inhibition with a threshold of *sc*_Ω_ and resorption factor R activation with a strength of *?*_Ω_ and a threshold of *r*_Ω_. Bone density decreases through bone resorption by osteoclasts at a rate *λ*_C_. To determine BMD, we scale bone density by bone mineral content, which is a constant BMC_0_, so we have BMD = BMC_0_Bd.

To summarize, our model extensions introduce three new parameters fit to data: the factor for peak increase in osteocyte apoptosis due to surgery (*η*_surg_), the factor for peak increase in osteoclast differentiation due to surgery (*ω*_surg_), and the timescale during which these effects last (*τ* ). The model parameters and species definitions for natural menopause are listed in Table 1, the initial conditions for the variables are listed in Table 2, and the model parameters and species definitions for surgical menopause are listed in Table 3.

**Table 1.**
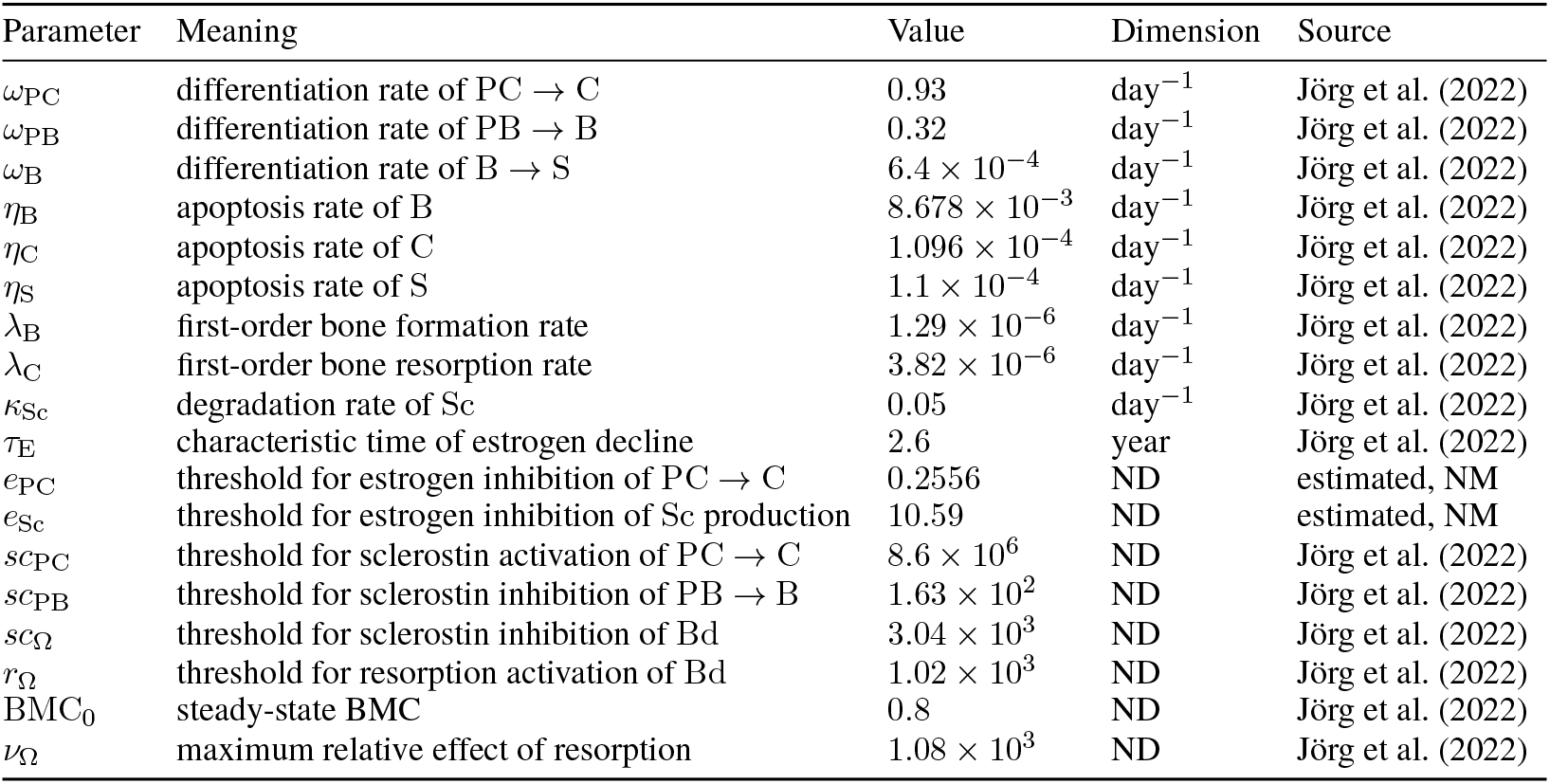
Natural menopause (NM) model parameters with parameters taken from Jörg et al. (2022) or estimated using the procedures outlined in Section 2.3. ND: no dimensions.

**Table 2.** Initial conditions taken at steady state at 30 years before menopause onset. All variables are in dimensionless form.

| Variable | Meaning | Initial value |
| --- | --- | --- |
| PB | preosteoblasts | $2.16 \times 10^2$ |
| PC | preosteoclasts | $4.10 \times 10^3$ |
| C | osteoclasts | $4.20 \times 10^1$ |
| B | osteoblasts | $1.07 \times 10^2$ |
| S | osteocytes | $6.12 \times 10^2$ |
| Sc | sclerostin | $1.12 \times 10^4$ |
| Bd | bone density | 1 |

**Table 3.** Surgical menopause (SM) model parameters from sources, calculated as described in Section 2.2, and estimated using the procedures outlined in Section 2.3 for short-term (15 years or less post-surgery) and long-term (up to 30 years post-surgery) data. ND: no dimensions.

| Parameter | Meaning | Value | Dimension | Source |
| --- | --- | --- | --- | --- |
| $\kappa_E$ | estrogen degradation rate post-surgery | 6.1996 | $\text{day}^{-1}$ | Ginsburg et al. (1998) |
| $k_{\text{syn}}$ | estrogen synthesis rate post-surgery | 0.6 | $\text{day}^{-1}$ | calculated |
| $E_{\text{surg}}(t_m + 30)$ | estrogen concentration 30 days post-surgery | 0.096 | ND | Bellanti et al. (2013) |
| $\eta_{\text{surg}}$ | apoptosis of S post-surgery | 5 | ND | estimated, SM short-term |
| $\tau$ | timescale of post-surgery dynamics | $9.7 \times 10^{-3}$ | $\text{day}^{-1}$ | estimated, SM short-term |
| $\omega_{\text{surg}}$ | enhanced C differentiation post-surgery | 1.86 | ND | estimated, SM short-term |
| $\eta_{\text{surg}}$ | apoptosis of S post-surgery | 0.4174 | ND | estimated, SM long-term |
| $\tau$ | timescale of post-surgery dynamics | 0 | $\text{day}^{-1}$ | estimated, SM long-term |
| $\omega_{\text{surg}}$ | enhanced C differentiation post-surgery | 0.2155 | ND | estimated, SM long-term |

### 2.3 Model solution and parameter estimation

We are interested in understanding and parameterizing the impacts of natural and surgical menopause on the dynamics of bone mineral density, Bd. Therefore, we solve the model ODEs and algebraic equations Eqs. (1)–(12) using `ode45` in MATLAB, with absolute and relative tolerances as 10^*−*8^, from 30 years before menopause onset to 30 years after menopause onset. Thus, the initial condition is 30 years before menopause onset for either natural or surgical menopause. For each simulation, we initialize the dynamic species based on their steady-state values at the initial condition. The Jörg et al. (2022) model includes premenopausal BMD decline. To determine the steady state solutions to Eqs. (4), (5), (7)–(9), and (11) while Bd is fixed at a value of 1, we solve the system of equations with time derivatives set to 0 using the `fsolve` function in MATLAB. The other species do not depend on Bd. This initialization is called inside the parameter estimation routine for the natural menopause case as the fitted parameters do affect the initial conditions. These updated values are reported in Table 2.

To estimate parameters in both the natural and surgical menopause mechanisms, we use the `lsqnonlin` function in MATLAB, which solves the following nonlinear least-square objective function using the Levenberg-Marquardt algorithm (Levenberg, 1944; Marquardt, 1963):

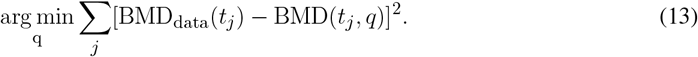

Here, BMD_data_(*t*_*j*_) is the natural BMD data at measurement times *t*_*j*_, and BMD(*t*_*j*_, *q*) is the BMD predicted by the model at the same times using parameters *q*. For each parameter estimation procedure, `lsqnonlin` algorithm options are set to tolerances of 10^*−*8^ and a maximum of 10,000 function evaluations and iterations. We use MATLAB version R2024b in Windows on a PC with 11th Generation Intel Core i7-11700 8 Core processor.

With the natural menopause BMD data described in Section 2.1, we first aim to re-estimate model parameters related to the impact of estrogen on osteoclastogenesis and the production of sclerostin by osteocytes for natural menopause. This corresponds to parameters *q*_NM_ = *{e*_PC_, *e*_Sc_*}*, and the estimated values of these parameters are listed in Table 1. With the estimated *q*_NM_, we then use surgical menopause BMD data to estimate surgical menopause parameters *q*_SM_ = {*η*_surg_, *τ, ω*_surg_} in the 15 years (short-term) and in the 30 years (long-term) after onset of menopause due to surgery. For the surgical menopause cases, upper and lower bounds are used to constrain parameter space. The lower bounds are *q*_SM_ = 0, 0, 0, which signify no effect, permanent effect, and no effect, respectively. The upper bounds are more subjective and were selected to yield only reasonable responses; we would need cell population data to further interrogate these with different upper bounds. The upper bounds used are *q*_SM_ = {5, 1*/*365, 5}, signifying that the peaks of *η*(*t*) and *ω*(*t*) at menopause onset are 5+1 = 6 times higher than baseline values in natural menopause and the surgery induced effects last a minimum of 1 day. The estimated values of these parameters are listed in Table 3.

We calculate sensitivity by varying the surgical menopause model parameters *q*_SM_ = {*η*_surg_, *τ, ω*_surg_} by 25% relative to the best-fit parameters. For instance, the upper bound is calculated using a 25% increase in *η*_surg_ and *ω*_surg_ and a 25% decrease in *τ*, which corresponds to longer-lasting effects of surgery.

## 3 RESULTS

With the mathematical framework introduced in Section 2.2, we study how mechanisms mediated by estrogen loss impact the bone remodeling process in both natural and surgical menopause. We focus on how BMD is affected decades after menopause and compare the BMD dynamics to both natural and surgical menopause data described in Section 2.1. We use data from natural and surgical menopause patients (Figure 1) and the mathematical models outlined in Section 2.2 with the methods described in Section 2.3 to parameterize and compare the model results with the relevant data. With new parameters and mechanisms, we show which important pathways impact BMD decline and rebound and propose new treatment directions based on our results.

### 3.1 Parameterized model of natural menopause captures BMD behavior in larger dataset

To assess the model output, we compare the BMD from the model with experimental data described in Section 2.1. Note that model parameters in Jörg et al. (2022) were estimated from natural menopause data in Looker et al. (1998), where only two BMD values were recorded after menopause; furthermore, these data were aggregated based on age groups and did not specify menopause onset, which can lead to underestimates in BMD loss. In Figure 3a for the natural menopause case, we present the average BMD for natural menopause patients in red markers for the larger dataset and distinguish data (Looker et al., 1998) that were used to parameterize the Jörg et al. (2022) model mathematical model with open red markers. The Jörg et al. (2022) model results, shown as the red dashed curve, fit the Looker et al. (1998) data well but do not fit the additional natural menopause data, which show a faster BMD decline.

**Figure 3.**
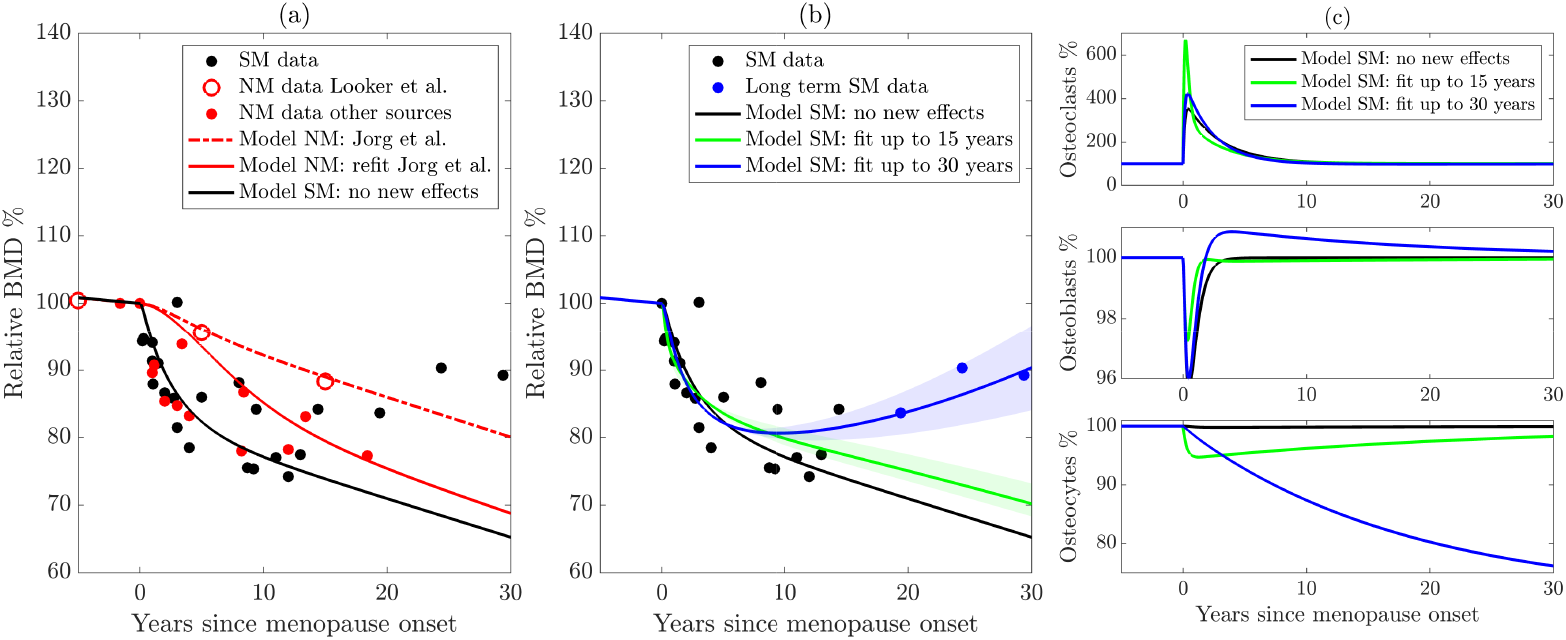
(a) Reparameterization of the Jörg et al. (2022) model using more sources of natural menopause (NM) data. The original model is shown by the dashed red curve, the new parameterization is shown by the solid red curve, and the black solid curve shows the model with sudden estrogen loss alone, without any new effects. (b) Extension of the model to surgical menopause. New effects are parameterized using long-term (blue curve) and short-term data (green curve). The shaded regions highlight model sensitivity to parameters. (c) Cell populations from the surgical menopause model with the three parameter sets from panel (b). NM: Natural menopause. SM: surgical menopause.

To improve the accuracy of the model after menopause, we estimate key parameters involved in natural menopause using the aggregated natural menopause data (Figure 1a). In particular, we estimate functional thresholds that depend on estrogen in both preosteoclast to osteoclast differentiation and sclerostin production from osteocytes, which are *e*_PC_ and *e*_Sc_, respectively. Following estimation of *q*_NM_ = {*e*_PC_, *e*_Sc_} for natural menopause (Table 1), we compare the relative BMD over time with natural menopause data in Figure 3a. Our reparameterized model of natural menopause shown as a red solid curve captures this rapid BMD decrease, and this behavior is within the experimental error (Figure 1a).

### 3.2 With new mechanisms, model of surgical menopause reproduces BMD trends in both short- and long-term data

Figure 3b shows the average relative BMD of surgical menopause patients 15 years post-surgery by black markers and long-term data beyond 15 years by blue markers, which indicate a rebound in BMD for these patients. Note that these are the same data in Figure 1b, where the error bars are shown. To illustrate that new mechanisms are necessary to capture the surgical menopause behavior, we first show the model dynamics (Figure 3a,b black curve) using newly estimated bone parameters for natural menopause from Section 3.1 and the estrogen decline in surgical menopause from Eq. (3) without including any new effects on cell dynamics, i.e., *η*_surg_ = 0 and *ω*_surg_ = 0. The effect of sudden estrogen loss alone enhances BMD loss compared to natural estrogen decline in Figure 3a. Indeed, the model with no new cell dynamics closely matches the surgical menopause data for the first 1–3 years post-surgery but overpredicts the extent of BMD loss in later years (Figure 3a,b black curve). This result suggests that surgical menopause involves both an increased loss of bone in the short term and a subsequent slowing of bone loss in the long term, which is not captured by simply including the onset of sudden estrogen decline in the natural menopause model by Jörg et al. (2022).

We estimate the new parameters modulated by surgical menopause in our model (Table 3): the percentage increase osteocyte apoptosis rate (*η*_surg_), the increased differentiation rate of preosteoclasts (*ω*_surg_), and the timescale over which the effects occur (*τ* ), using data in Figure 1b. We use the parameterization methods in Section 2.3 with *q*_SM_ = {*η*_surg_, *ω*_surg_, *τ*} . To investigate the differences in BMD over both “short” and “long” time scales, we fit the surgical menopause model to surgical menopause data up to 15 years and again up to 30 years to assess parameter differences (Table 3). We obtain root mean squared errors in BMD prediction of 4.59% and 4.63% compared to the surgical menopause data up to 15 years and 30 years, respectively. Using the short and long time scale datasets resulted in different long-term BMD dynamics (Figure 3b). The short-term surgical menopause model better captures the data for the 15-year period compared to the case without new effects (Figure 3a,b black curve). The long-term surgical menopause model also fits these data well and has a BMD rebound not observed in the short-term case. The sensitivity of the model predictions to the parameter values is shown by the corresponding shaded area around each curve. From these sensitivity regions, the behavior of the model in the 2 years post-surgery is not sensitive to the new effects, but the long-term predictions of the model are altered substantially by parameter variations.

To understand other differences in these parameterized surgical menopause models, we simulated the dynamics of the osteoclast, osteoblast, and osteocyte cell populations after menopause (Figure 3c). The no new effects case and the short-term case show that all species return to approximately their premenopausal levels within 10 years. This is expected, as the timescale of surgical effects, *τ*, is small for the short-term model. The short-term effect is seen in the osteoclast (osteocyte) population, where a sharp increase (decrease) is seen at the onset of menopause and then rebounds. This sudden increase in osteoclast population yields steeper dips in Figure 3b, i.e., relative BMD % in the short-term surgical menopause model is lower than other models immediately after menopause onset.

In the long-term surgical menopause case, we see a smaller increase in osteoclastogenesis after menopause onset compared to the short-term case; *ω*_surg_ is an order of magnitude smaller in the long-term case than the short-term case. Osteocyte density also decreases more slowly in the long-term surgical menopause model, but continues after the onset of menopause; *τ* = 0 for long-term surgical menopause effects making the impact of surgery permanent. Interestingly, the osteoblast percentage decreases after surgery, then rebounds to a slightly higher level for an extended period of time. The long-term surgical menopause model yields lower and continually declining osteocyte populations and higher but leveling off osteoblast populations. Since sclerostin is downregluated and bone formation is upregulated, this results in an increase in BMD over long periods of time, enabling the rebound in BMD.

## 4 DISCUSSION

We present a mathematical model of the bone remodeling process to understand the effects of estrogen loss due to surgical menopause. Since experimental data suggest that surgical menopause and natural menopause impact different mechanisms in bone remodeling, we extended an existing mathematical model of bone remodeling to incorporate the increased differentiation mechanisms of osteoclasts and increased apoptosis of osteocytes. The objective of this framework is to understand and capture trends seen in newly aggregated physiological data: (1) surgical menopause leads to an increased loss of BMD in the short term, and (2) this loss slows or even rebounds by 10 or more years post-surgery.

The BMD predictions after reparameterization to the larger natural menopause BMD dataset better capture the overall trends of the natural menopause data compared to the previous mathematical model (Jörg et al., 2022). These parameters, *e*_PC_ and *e*_Sc_, influence the strength of estrogen signaling on osteoclastogenesis and sclerostin release by osteocytes. Compared to the previous mathematical model, our parameter fitting results in smaller threshold values for the estrogen signaling pathway related to osteoclastogenesis, resulting in more osteoclast differentiation for a higher concentration of estrogen. These parameter changes capture the overall larger decrease in BMD shown in the newly aggregated data.

Our new surgical menopause model that incorporates mechanisms impacted by the sudden loss of estrogen and inflammation resulting from surgery, including increases in osteocyte apoptosis and osteoclastogenesis follows the sharp decrease in BMD in the first 15 years post-surgery and a rebound in BMD 15–30 years post-surgery, consistent with the clinical data. To understand what mechanisms play a role in this varied behavior, we fit our surgical menopause model to two datasets: BMD data up to 15 years and up to 30 years post-surgery. The short-term data fit indicates a higher apoptosis rate of osteocytes and a lower differentiation rate of preosteoclasts compared to the long-term data fit. Osteocyte levels differ substantially between the long-term and short-term model calibrations with the long-term parameters yielding slower but permanent rates of osteocyte apoptosis due to surgical menopause.

Our mathematical study does not include hormonal interventions or bone remodeling treatments. Most osteoporosis treatments are classified as anti-resorptive agents that inhibit resorption by osteoclasts to prevent further bone loss but can cause side effects. For example, bisphosphonates are one of the most widely prescribed treatments (Khosla and Hofbauer, 2017) to promote osteoclast apoptosis (Berkhout et al., 2015), but they can lose efficacy over time and lead to osteonecrosis and atypical femoral fractures (Shane et al., 2014; Khosla et al., 2007; Whitaker et al., 2012). Another anti-resorptive treatment is the anti-RANKL monoclonal antibody denosumab, which inhibits osteoclastogenesis (Khosla and Hofbauer, 2017). New antibody treatments targeting sclerostin production from osteocytes show promising results that increase bone formation and decrease bone resorption (Recker et al., 2015; Zhang et al., 2016; Chavassieux et al., 2019), but bone formation only lasts a few months (Ominsky et al., 2017) and therefore treatment is only recommended for 6–12 months (McClung, 2017). Our parameter estimates indicate promising avenues of treatment for patients undergoing surgical menopause, where our results suggest treatments targeting osteocyte or osteoblast dynamics may support long-term BMD preservation. Implementing treatment methods on this system requires further investigation.

## CODE AVAILABILITY

We have provided model code and files in a repository at https://github.com/ashleefv/SurgicalMenopauseBone (Nelson et al., 2025).

## AUTHOR CONTRIBUTIONS

- A.C.N: Conceptualization, data curation, formal analysis, investigation, methodology, software, validation, visualization, writing–original draft, writing–review and editing.
- E.F.Y.: Conceptualization, data curation, formal analysis, investigation, software, validation, visualization, writing–original draft, writing–review and editing.
- Y.Z.: investigation, methodology, writing–review and editing.
- C.V.C.: Conceptualization, investigation, methodology, software, visualization, writing–original draft, writing–review and editing.
- S.F.H.: data curation, investigation, methodology, writing–review and editing.
- L.K.B.: investigation, methodology, writing–original draft, writing–review and editing.
- P.D.: investigation, methodology, writing–review and editing.
- S.G.: investigation, methodology, writing–review and editing.
- B.J.S.: Conceptualization, funding acquisition, writing–review and editing.
- A.N.F.V: Conceptualization, formal analysis, funding acquisition, investigation, methodology, project administration, software, supervision, visualization, writing–review and editing.

## ACKNOWLEDGMENTS

This work resulted from the “Sex Differences in Physiology: Mathematical Modelling and Analysis” workshop at the Banff International Research Station (BIRS) on March 5–10, 2023. We would like to thank the staff and the organizers for convening the workshop and other participants in the workshop for helpful conversations and feedback related to this project.

